# Retrograde viral delivery of hyper-IL-6 activates Stat3 in corticospinal tract neurons but causes severe tremors and weight loss in adult mice

**DOI:** 10.64898/2025.12.30.697090

**Authors:** Zimei Wang, Pantelis Tsoulfas, Murray G. Blackmore

## Abstract

Cytokines in the IL-6 family can improve axon regeneration after nervous system injury, acting in part by stimulating Stat3 phosphorylation and pro-regenerative signaling in injured neurons. Hyper-IL-6 (hIL-6) is an engineered ligand with enhanced signaling properties, shown previously to stimulate axon growth even more effectively than native IL-6 family members. Here we tested a method of hIL-6 delivery based on injection of retrograde AAV to the cervical spinal cord of adult mice. This was envisioned as a first step toward a regenerative treatment after spinal cord injury based on widespread stimulation of descending projection neurons. We found, however, that animals treated with AAV2-retro-hIL-6 developed severe tremors and weight loss within days of injection and reached criteria for humane euthanasia within five to eleven days, depending on viral dose. Examination of the cortex showed a pronounced increase in Stat3 phosphorylation in corticospinal tract (CST) neurons in hIL-6 animals versus control, confirming effective retrograde signaling. Stat3 phosphorylation and evidence of microglial activation were detected in tissue surrounding CST cell bodies and along descending CST axons, revealing extensive cytokine signaling and microglial response associated with secreted hIL-6. These results confirm the ability of hIL-6 expression to activate Stat3 signaling in CST neurons but uncover the potential for severe negative effects that must inform any further development of hIL-6 as a pro-regenerative therapeutic.

**SIGNIFICANCE STATEMENT:** There is an urgent need to develop therapeutic strategies to improve repair after nervous system injury. In this study, we tested a new way to deliver Hyper-IL-6, an engineered cytokine with enhanced signaling properties, based on retrograde adeno-associated injection to the spinal cord and widespread uptake by descending axons. Expression of hIL-6 activated an expected pathway in descending neurons but also caused severe side effects and signs of widespread microglial activation in the cortex. These outcomes confirm potent signal activation by hIL-6 but reveal severe off-target effects that must be considered in any future application as a pro-regenerative treatment.

## INTRODUCTION

A central goal in regenerative neuroscience is to develop effective molecular strategies to improve axon growth after damage to the central nervous system (CNS), especially for long-distance projection neurons that connect the brain and spinal cord. Axon growth depends in part on ligand/receptor interactions that trigger pro-regenerative signaling cascades in the neuron (He and Jin, 2016; Tedeschi and Bradke, 2017; Vartak et al., 2023). Thus, identifying and delivering such regeneration promoting signaling molecules is a prominent strategy to enhance CNS repair (O’Donovan, 2016; Mahar and Cavalli, 2018).

Cytokines in the IL-6 family have emerged as a leading class of pro-regenerative ligands. The family is characterized by the use of the signaling GP130 receptor, in association with ligand-specific co-receptors, leading primarily to activation of JAK/STAT signaling, with contributions from additional pathways such as PI3K and MAPK (Zigmond, 2012; Rose-John, 2018). In the peripheral nervous system, where axon regeneration occurs spontaneously, IL-6 family members are upregulated at sites of injury, retrogradely transported to neuronal cell bodies, and contribute to the regenerative response (Zhong et al., 1999; Cafferty et al., 2004; Cao et al., 2006). Extensive prior work has demonstrated IL-6 family members delivered by repeated injection or viral expression promote axon regeneration in injured retinal ganglion cells, sensory neurons, sympathetic neurons, and motor neurons (Sahenk et al., 1994; Cui and Harvey, 2000; Cao et al., 2006; Leaver et al., 2006a; Smith et al., 2009; Sun et al., 2011; Zigmond, 2012). However, manipulation of IL-6 signaling has produced more modest and mixed effects on axon regeneration in corticospinal tract (CST) neurons, important mediators of descending sensorimotor control. A variety of molecular strategies have been tested in CST neurons, including administration of IL-6 ligands (Blesch et al., 1999; Yang et al., 2012, 2015; Hodgetts et al., 2018), direct expression of Stat3 (Lang et al., 2013), and signal amplification by deletion of Socs3, a feedback inhibitor of Jak/Stat signaling (Jin et al., 2015; Geoffroy et al., 2022). Although these approaches have yielded pronounced long-distance growth in the optic and sensory systems, their effects in CST remain limited to short-distance arborization near spinal injury sites (Blesch et al., 1999; Hodgetts et al., 2018; Yang et al., 2012), and show lower efficacy when used in fully mature animals or when applied after injury (Jin et al., 2015; Geoffroy et al., 2022).

How can growth stimulation by IL-6 cytokines be enhanced, especially in relatively unresponsive neuronal populations such as CST? One promising strategy involves hyper-IL-6 (hIL-6), an engineered cytokine in which IL-6 is pre-linked to the functional region of the IL-6Rα receptor (Fischer et al., 1997). The key advantage of this approach is that bypasses reliance on ligand-specific co-receptors, whose low availability is rate limiting, and instead signals directly through GP130 receptors, which are ubiquitously expressed across most cell types, including CST neurons (Zigmond, 2012; Jin et al., 2015; Wang et al., 2025). This direct engagement results in robust and sustained JAK/STAT signaling in recipient cells (Fischer et al., 1997; Rose-John, 2012, 2018; Fischer, 2017). Indeed, in retinal ganglion cells, axon regeneration from viral expression of hIL-6 greatly exceeded the effects of CNTF or IL-6 (Leaver et al., 2006b; Leibinger et al., 2013, 2016). Furthermore, viral delivery of hIL-6 by focal injection to the cortex enhanced CST axon growth and axon regeneration by subcortical neuronal populations that received hIL-6 by secondary transsynaptic secretion (Leibinger et al., 2021).

We hypothesized that further gains might be achieved if hIL-6 delivery could be made more widespread throughout the CST and other descending neuronal populations. We expressed hIL-6 in adult mice via retrograde injection of AAV2-retro to cervical spinal cord, shown previously to transduce most CST neurons and tens of thousands of additional supraspinal neurons in midbrain and brainstem (Wang et al., 2018, 2022; Beine et al., 2022). We reasoned that retrograde transduction would lead to hIL-6 secretion and autocrine activation of Jak/Stat signaling throughout the supraspinal connectome, potentially elevating regenerative ability in numerous populations. We found, however, that retrograde hIL-6 resulted in a rapid and severe decline in health, with animals developing pronounced tremors and weight loss within days of AAV injection. Examination of the cortex confirmed elevation of pStat3 in CST cell nuclei but also extensive pStat3 signal and microglial activation in surrounding tissue. These observations confirm the ability of hIL-6 to stimulate Jak/Stat signaling in CNS cells while highlighting negative whole-body consequences that caution against excessive expression, and which may limit hIL-6’s practical application as a pro-regenerative stimulus.

## METHODS

### Animal details

Male and female C57BL/6J mice (8–12 weeks old, The Jackson Laboratory) were used in all experiments. Animal procedures were carried out in full compliance with the National Institutes of Health *Guide for the Care and Use of Laboratory Animals* and all protocols were reviewed and approved by the Marquette University Institutional Animal Care and Use Committee (IACUC protocol numbers 3283 and 4013). No sex-related differences were detected. Mice were housed with unrestricted access to food and water under a controlled 12 h light/12 h dark cycle, with temperature maintained at 22 ± 2 °C and relative humidity at 40–60%.

### Plasmid construction and AAV preparation

rAAV2-retro-CAG-H2B-mGreenLantern (Addgene#177332) was produced at the University of Miami viral core facility at the Miami Project to Cure Paralysis, titer = 1.4x10^13^ particles/ml. rAAV2-retro-tdTomato (Addgene #59462) were produced at the University of North Carolina Viral Vector Core at 3x10^12^ particles/ml. The coding sequence for hyper-IL-6 was created following the design of (Fischer et al., 1997), with sequences 113-323 of the soluble IL-6R attached to sequences 29 to 212 of human IL-6 by a RGGGGSGGGGSVE linker. The resulting sequence was codon-optimized using a Genscript algorithm for mouse expression and synthesized by Genscript with flanking KpnI and EcoRI restriction sites. The resulting construct was inserted into an AAV construct with a CAG promoter, using (Addgene# 191093). rAAV2-retro-Hyper-IL-6 was then produced at the University of Miami viral core at a titer of 2.1x10^14^. Prior to injection all viruses were diluted in Phosphate Buffered Saline (PBS: Sigma P4474, 154mM sodium chloride, 1.058mM potassium phosphate monobasic, 2.97mM sodium phosphate dibasic dihydrate, pH 7.4) with 5% D-Sorbitol (Sigma S1876) and 0.001% Poloxamer 188 Surfactant (ThermoFisher 24040032). Final titers in the injected mix of AAV were rAAV2-retro-H2B-mGl (1.0x10^12^ ), rAAV2-retro-CAG-tdTomato (1.0x10^12^ ), and rAAV2-retro-hIL-6 (2.5x10^12^ or 1.0x10^13^). In control animals, AAV2-retro-Malat-Barcode6 ((Wang et al., 2025) was substituted for AAV2-retro-hIL-6 at a titer of 1.0x10^13^.

### Surgical procedures

All surgical procedures were conducted under ketamine/xylazine anesthesia in adult C57BL/6 mice (8–12 weeks old; 20–28 g). For AAV delivery, animals were secured in a custom spinal stabilization device, and a laminectomy was performed at the C5 level. AAV-retro vectors were administered using a 1701 Hamilton syringe attached to a pulled glass capillary needle and controlled by a Stoelting QSI pump (catalog #53311) mounted on a micromanipulator. Virus was infused at 0.04 μl/min, targeting a site 0.35 mm lateral to the midline. The injection was performed first at a depth of 0.8 mm (500 nl), after which the needle was retracted to 0.6 mm and the remaining 500 nl was delivered.

### Tissue Processing and Imaging

Animals were euthanized by CO_2_inhalation and immediately underwent transcardial perfusion with 0.9% saline and 4% paraformaldehyde (PFA) solutions in 1× phosphate-buffered saline (PBS) (15710, Electron Microscopy Sciences, Hatfield, PA). Brains were then removed and post-fixed overnight in 4% PFA at 4°C, embedded in 12% gelatin in PBS and cut via Vibratome to yield 100 μm sections. Prior to pStat3 detection, sections underwent an antigen retrieval procedure of immersion in a solution of 40mM Citrate (J63950.AP, Thermo Scientific) in water for thirty minutes at 80°C followed by rinsing in PBS. Sections were incubated overnight with primary antibodies pSTAT3 (Cell Signaling 9145 1:500, RRID:AB_2491009) or Iba1 (WAKO 019-19741 1:500, RRID: AB_839504).

Sections were then rinsed and incubated for 2 hours with Alexa Fluor 647-conjugated secondary antibodies (ThermoFisher, Waltham, MA, 1:500, A-21245, RRID AB_2535813). Fluorescent images were acquired using a Nikon A1R+ laser scanning confocal microscopy system on a Nikon Ti2-E inverted microscope. For quantification of cJUN and ATF3, regions of interest (ROIs) were first manually drawn in NIS Elements software using the mScarlet label within retrogradely transduced CST cell bodies. The average pixel intensity of the detection channel was then determined for each ROI. All visible CST cell bodies from at least two replicate cortical sections for each animal were selected by observers blinded to treatment.

### Experimental design and statistical analyses

For quantification of immunofluorescence experimental replicates are indicated in figure legends and significant differences were tested by one way ANOVA with post-hoc Dunnett’s using Prism software (GraphPad version 10.2.3). Statistical details and replicate numbers for each experiment can be found in the figure legends. All quantification of images was conducted by experimenters blind to experimental group.

## RESULTS

DNA coding for the open reading frame of hIL6 was synthesized and packaged in AAV2-retro (**Fig. 1A**). Prior to injection, AAV2-retro-hIL6 was mixed with AAV2-retro-H2B-mGreenLantern (mGl), a nuclear-localized fluorescent reporter, and AAV2-retro-tdTomato, a cytoplasmic reporter. Final titers of AAV2-retro-hIL6 were either 1x10^13^ particles per ml (hereafter high titer) or 2.5x10^12^ particles per ml (hereafter mid titer). Control animals received AAV2-retro-H2B-mGl along with control AAV2-retro-Malat-Barcode at 1x10^13^ particles per ml (Wang et al., 2025) to equalize total viral load. Adult mice of both sexes received injection of virus to cervical spinal cord. We have shown in previous studies that this approach yields reliable retrograde transduction of tens of thousands of supraspinal projection neurons distributed throughout the brain, including about 8,000-10,000 CST neurons on each side of the cortex (Wang et al., 2018, 2022) . **Table 1** provided surgical details and viral titers for all animals used in this study.

**Figure 1.**
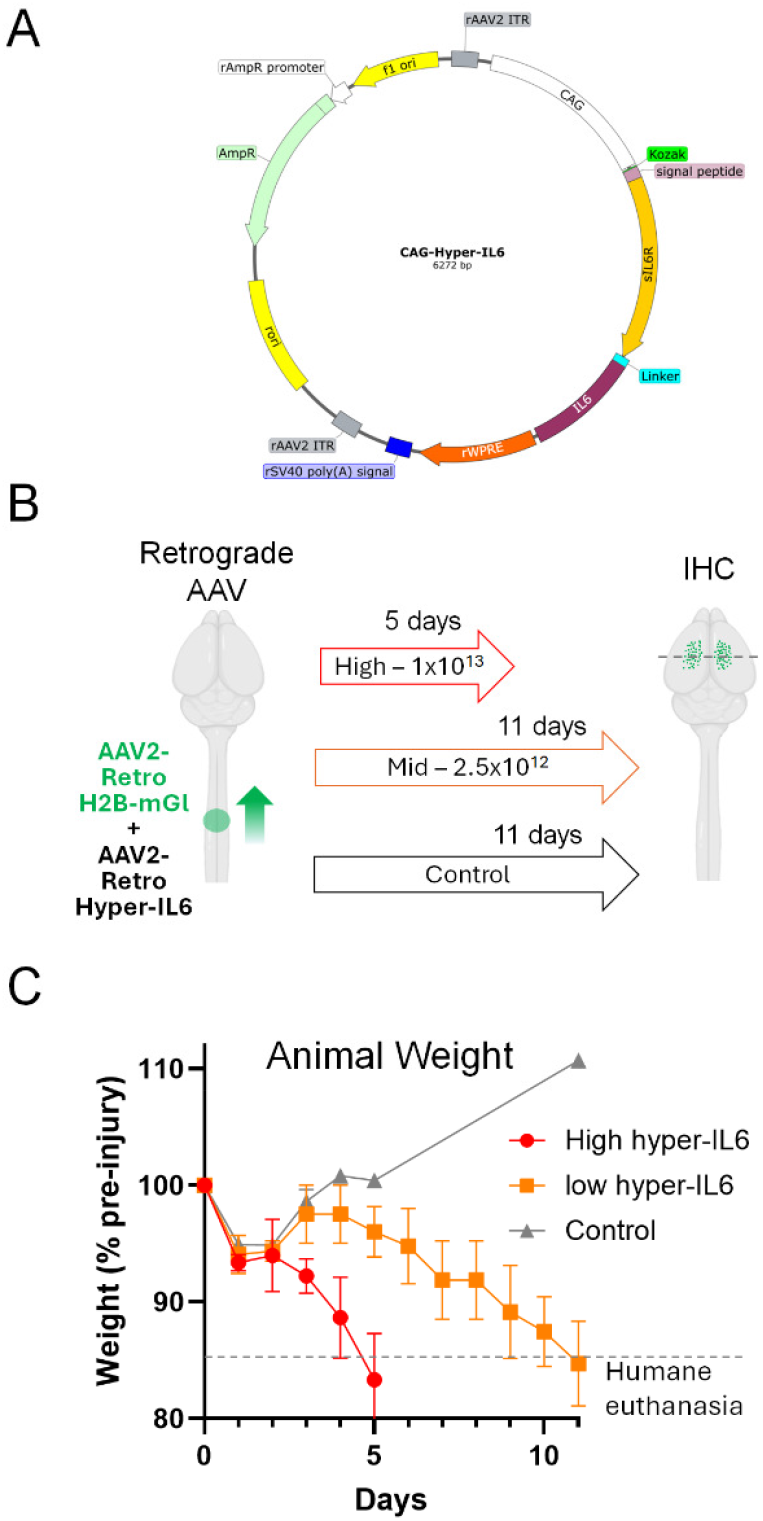
Retrograde viral delivery of hyper-IL6 from cervical spinal cord causes rapid weight loss in adult mice. (A) A map of the plasmid used to produce AAV2-Retro-Hyper-IL6. (B) Experimental overview showing retrograde viral delivery and the experimental timeline. The original plan of 14 days survival was shortened for reasons of animal welfare. (C) Animal weights normalized to preinjury levels. Control animals showed a transient postinjury decrease in weight followed by weight gain. Animals receiving AAV2-Retro-Hyper-IL6 displayed post-injury weight loss comparable to control followed by subsequent loss of weight. Animals were sacrificed when weight fell below pre-set 15% threshold set by animal well-being (dotted line). N= 3 animals per group.

Animal weights were tracked after surgery as an indicator of animal well-being and are plotted in **Figure 1C**. Control animals showed post-surgery weight loss of about 5%, which is typical for this procedure. Weights remained suppressed through the second day post-surgery, followed by recovery to pre-injury levels by day four. Animals that received hIL6 mirrored control animals for the first two days, with both the mid-titer and high-titer displaying a similar 5% reduction that stabilized between days one and two. The groups diverged after this, with the high-titer hIL-6 group undergoing further weight loss that by five days post-surgery reached more than 15% decline, the threshold for humane euthanasia set by local animal research oversight. In contrast, the mid-titer hIL6 showed weight gain between days two and three, similar to controls, but then a gradual decrease that reached the threshold of human euthanasia by eleven days post-surgery. Importantly, weight loss was accompanied by the development of severe tremors and decline in mobility, with animals observed to adopt a hunched posture with readily visible whole-body shaking. This phenotype developed between days three and four in the high-titer group and around day six in the mid-titer group. Thus, spinal injection of AAV2-retro-hIL6 to adult mice causes rapid weight loss and the development of whole-body tremors with dose-dependent onset and rate of progression.

We next examined cortices for evidence of hIL-6 activity, using phosphorylation of Stat3 (pStat3) as an indicator (Fischer et al., 1997; Leibinger et al., 2016, 2021). Animals were perfused, cortical sections prepared, and immunohistochemistry for pStat3 was performed. As expected, cell nuclei labeled with H2B-mGl were readily visible throughout layer five of cortex, marking the location of retrogradely transduced CST neurons. In control tissue signal for pStat3 was very low or absent, including in CST neurons (**Fig. 2A, D)**. In contrast, pStat3 signal was readily detectable in hIL-6 animals in both the mid-titer and high-titer groups (**Fig. B, C, E, F)**. The signal co-localized with CST neuronal nuclei, identified by H2B-mGl, (**Fig. 2E, F, arrows)** and quantitative analysis of nuclear pStat3 signal in cell nuclei revealed a strong and significant increase in hIL6-treated animals (Control: 3.61 ± 0.51 SEM, mid-titer hIL-6 30.29 ± 1.13 SEM, high-titer hIL-6, 23.22 ± 1.36 SEM p<0.0001 One Way ANOVA with post-hoc Dunnett’s) (**Fig. 2G**). pStat3 levels were slightly higher in the mid-titer group than the high-titer, which may reflect the longer survival time. In addition to signal in CST nuclei, pStat3 was also densely distributed throughout all cortical layers in both mid-titer and high-titer hIL-6 animals (**Fig. 2E, F**). This indicates that expression of hIL-6 in CST neurons leads not only to autocrine activation of Stat3 but also widespread paracrine activation of many cortical cells, likely via secretion. A similar phenomenon of transneuronal activation by secreted hIL-6 has been noted previously (Leibinger et al., 2021). In summary, retrograde transduction with hIL-6 vectors led to activation of Stat3 in transduced CST neurons and widespread Stat3 activation throughout cortical tissue.

**Figure 2.**
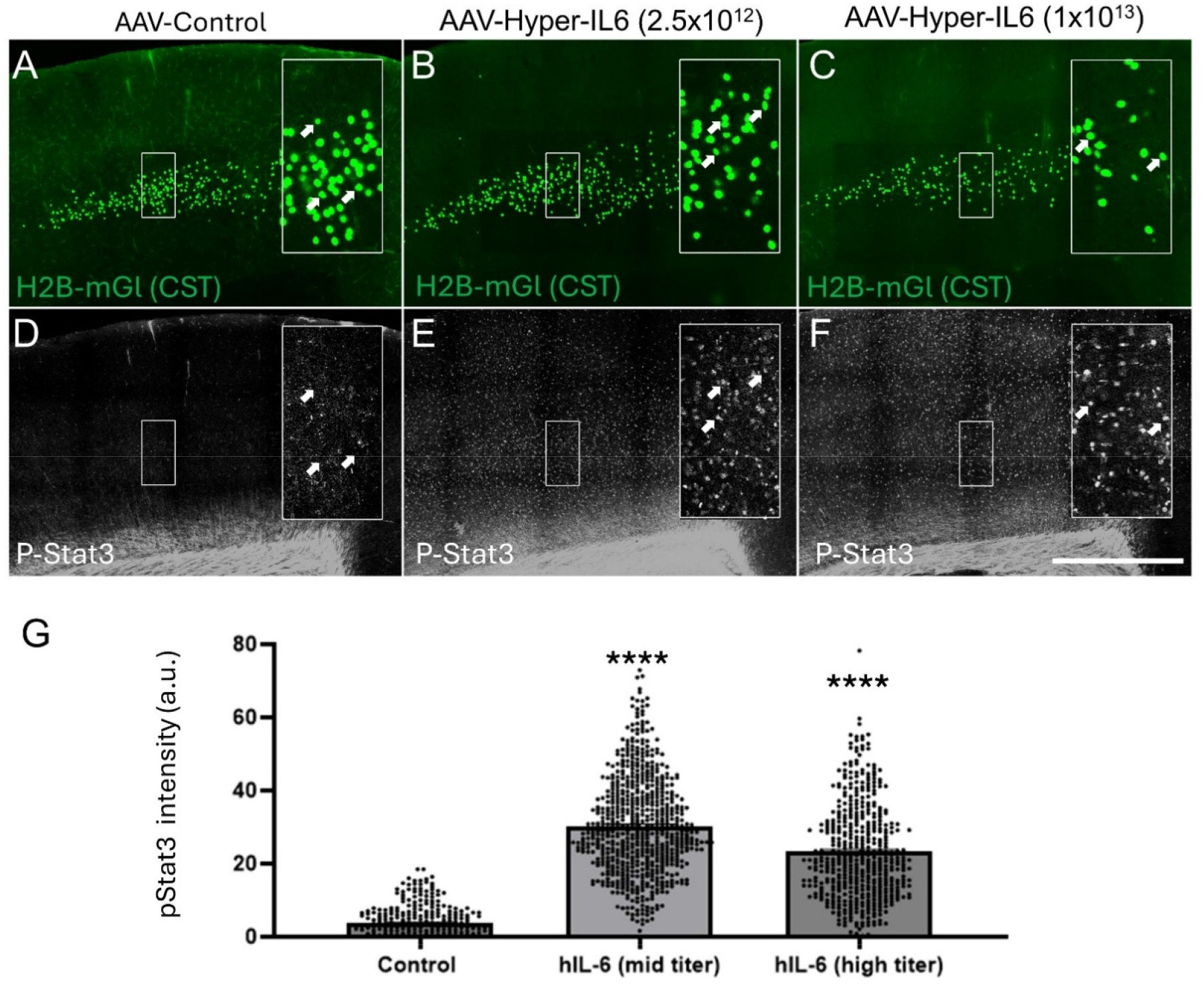
Retrograde viral delivery of hyper-IL6 leads to widespread Stat3 activation in cortical tissue. Adult mice received spinal injection of AAV2-Retro-H2B-mGI with AAV2-Retro-Malat-barcode control or with AAV2-Retro-Hyper-IL6 followed by immunohistochemistry for phospho-Stat3. (A-C) H2B-mGI (green) labels retrogradely labeled CST nuclei in cortex. (D-F) pStat3 signal is very dim in control (D) but readily detectable throughout the cortexes of animals that received Hyper-IL6 (E,F). Insets show CST cell nuclei (green) with co-localized pStat3 signal (arrows). Note that p-Stat3 signal also present in numerous surrounding nuclei, presumably reflecting paracrine activation by secreted hlL-6. (G) Quantification of signal intensity within individual CST nuclei, defined by regions of H2B-mGI signal, shows a significant increase in hlL-6 treated aniamls. ^****^ p<0.0001, One Way ANOVA with post-hoc Dunnett’s. N>100 nuclei from each of three animals per group. Scale bar is 0.5mm.

We then evaluated microglial activation as a component of the neuroinflammatory response in cortical tissue using immunohistochemistry for Iba1, a widely used marker of microglial reactivity. (Imai and Kohsaka, 2002). As previously, immunohistochemistry was performed on coronal sections of cortex, using H2B-mGl to identify CST cell nuclei (**Figure 3A-C**). In control animals, Iba1 signal was dimly visible in cells throughout the cortex, with a pattern of indistinct expression concentrated in the cell bodies (**Figure 3D**). In contrast, microglia exhibited an activated profile as evidenced by a markedly stronger Iba1 signal across all cortical layers in animals receiving mid-or high-titer hIL-6 displayed, with clear labeling of both microglial cell bodies and fine ramified processes (**Figure 3E,F**). Quantification confirmed a significant elevation of Iba1 signal intensity in hIL-6-treated animals compared to control (Control: 10.98 ± 0.46 SEM, mid-titer hIL-6 40.63 ± 2.95 SEM, high-titer hIL-6, 36.19 ± 1.50 SEM p<0.0001 One Way ANOVA with post-hoc Dunnett’s). In addition, by visualizing the cytoplasmic AAV2-retro-H2B-tdTomato reporter, it was revealed that Iba1 immunohistochemistry was concentrated in the region of CST cell bodies (**Figure 3H, Asterisk**) and also appeared along the track of descending axons (**Figure 3H, arrow**). Thus, retrograde delivery of AAV-hIL-6 to CST neurons caused extensive microglial with a spatial distribution that is consistent with CST neurons and axons acting as the source of activation. Taken in whole, our data support a model in which retrograde hIL-6 transduction of descending neurons leads to secretion and widespread elevation of Jak/Stat signaling and activation of microglial cells, accompanied by rapid-onset neurological dysfunction and weight loss.

**Figure 3.**
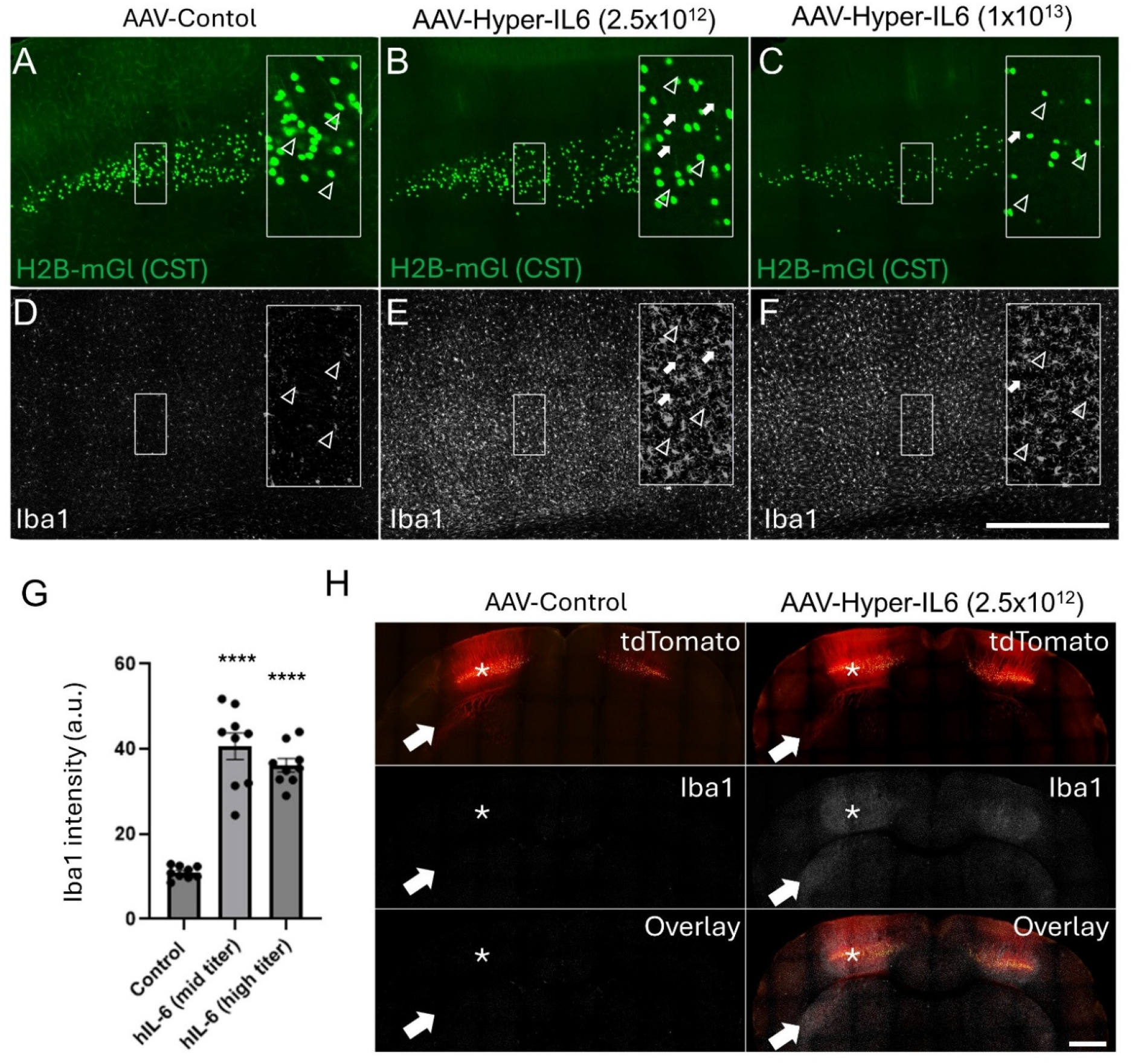
Retrograde viral delivery of hyper-lL6 leads to widespread microglial activation in cortical tissue. Adult mice received spinal injection of AAV2-Retro-H2B-mGI with control AAV2-Retro Malat-barcode or with AAV2-Retro-Hyper-lL6 followed by immunohistochemistry for lba1. (A-C) H2B mGI marks the location of retrogradely labeled CST nuclei in cortex. (D) lba1 signal is dim and appears confined to the main cell mass in animals that received only H2B-mGI (D). (E,F) lba1 signal is bright in cells throughout the cortex and extends into fine cellular processes, which indicates microglial activation. (G) Quantification of signal intensity in randomly selected layer V regions shows significant elevation of lba1 signal in hll6-treated animals (^****^ p<0.0001, one way ANOVA with post-hoc Dunnett’s). N=three randomly selected sample regions in each of three replicate animals per group. (H) Animals were spinally co-injected with AAV2-Retro-tdTomato and control AAV2-Retro-Malat barcode or AAV2-Retro-hll-6 followed by immunohistochemistry for lba1. lba1 signal is present in regions of CST cell bodies (asterisk) and descending axons (arrow) with hll-6 treatment (right) but not in controls (left). Scale bars are 0.5mm.

## DISCUSSION

We have tested a strategy of widespread hIL-6 transduction in descending neuronal populations, motivated by promising pro-regenerative findings in the optic system and experiments using focal injection of AAV-hIL-6 to the cortex (Fischer et al., 1997; Leibinger et al., 2016, 2021). Immunohistochemistry for pStat3 confirmed activation of Jak/Stat signaling in retrogradely transduced CST neurons but also found extensive pStat3 signal and microglial activation in neighboring cells. At the same time, animals rapidly developed signs of extreme nervous system dysfunction including whole-body tremors, lethargy, and loss of body weight. These findings confirm hIL-6’s effectiveness as a tool to trigger Stat3 activation in corticospinal tract (CST) neurons but also indicate that this strategy may be constrained by excessive hIL-6 signaling in off-target cells, including microglia, and rapidly emerging, severe side effects.

Prior evidence exists that overexpression of IL6 in the CNS can negatively affect animal well-being, for example from GFAP-IL-6 transgenic mice in which IL-6 secretion is increased in astrocytes (Campbell et al., 1993). Different founder lines displayed a range of IL-6 levels. Interestingly, the pups of lines with the highest IL6 expression displayed a phenotype highly reminiscent of our observations, with “hunched posture, piloerection, tremor, ataxia, hindlimb weakness, and seizures.” Affected lines also displayed reduced body size and died within weeks of birth. Thus, astrocytic expression of IL-6 seems to cause whole-body effects very similar to those of neural expression of hIL-6, with the main difference being a more rapid onset in the current study. We hypothesize this may reflect higher expression levels via AAV2-retro delivery and/or by the use of hyper-IL-6, which signals more potently than native IL-6 and which affects a wider range of cell types by depending solely on the ubiquitous GP130 receptor (Fischer et al., 1997; Rose-John, 2012). This precedent in the literature strengthens the conclusion that widespread secretion of IL-6 cytokine in the brain, especially the potent hIL-6 version, comes with a high risk of neurologic dysfunction.

Prior studies that delivered AAV-hIL-6 to the retina and cortex, however, did not report negative effects on animal health (Leibinger et al., 2016, 2021; Fischer, 2017). This may reflect a difference in distribution. Previous work employed focal injection of AAV-hIL-6 to the cortex, leading to secretion from cells near the site of injection and also anterograde secretion from axon terminals. In contrast, our prior work has established that cervical injection of AAV2-retro results in transduction of spinal cells at the site of injection, propriospinal neurons that project to cervical spinal cord, and tens of thousands of descending neurons distributed across dozens of populations in the brainstem, midbrain, and cortex (Wang et al., 2018, 2022; Beine et al., 2022). This retrograde distribution likely led to hIL-6 secretion with excessive cytokine signaling and microglial activation in numerous locations throughout the neuroaxis beyond the cortex. The present study focused on the cortex to demonstrate the ability of retrograde hIL-6 transduction to evoke tissue-level Jak/Stat and microglial activation. It remains unknown, however, whether cortical cytokine overexposure contributed to the tremors and other whole-body effects. Widespread or sustained cytokine exposure within the CNS most likely engages non-neuronal targets, including microglia, astrocytes, and vascular cells, leading to pronounced glial reactivity and amplification of inflammatory signaling. Enhanced microglial activation, as reflected by increased Iba1 immunoreactivity across cortical layers, likely reflects a state of heightened surveillance and reactivity that, if prolonged, will compromise synaptic integrity, circuit stability, and tissue homeostasis.

It is possible that the harmful effects of hIL-6 can be mitigated. Indeed, as suggested by prior work using focal injection, a more spatially restricted mode of delivery can likely avoid the extreme whole-animal malaise observed here. Indeed, if the harmful effects of hIL-6 were found to originate only from expression in restricted brain regions, it is possible that other regions could be found in which hIL-6 can be expressed without negative effects. In parallel, temporal control of hIL-6 expression, for example through inducible gene systems, could be used to terminate hIL-6 before the emergence of harm.

Therefore, by managing spatial and temporal distribution, experiments can likely still be performed that capitalize on hIL-6’s potent ability to activate Jak/Stat signaling in CNS cells while avoiding negative whole-animal effects.

From a translational perspective, our findings identify a key challenge for deploying hIL-6 as pre-regenerative therapy. While hIL-6 does effectively activates Stat3 in CST neurons, it also seems to diffuse and engage multiple surrounding cell types. Such widespread cytokine exposure raises concerns, as excessive cytokine signaling is known to produce detrimental long term effects. Thus, a broader implication of these findings may be that there are inherent risks to any strategy that relies on a secreted factor like hIL-6 to achieve Jak/Stat activation in the CNS, as this approach come with unavoidable activation of off-target cells and the potential to trigger of inflammatory feed-forward damage. Thus, molecular strategies that restrict Stat3 activation, ideally in a cell-autonomous manner that targets Jak/Stat activation specifically to damaged neurons, may be preferable.

## REFERENCES

Beine Z, Wang Z, Tsoulfas P, Blackmore MG (2022) Single nuclei analyses reveal transcriptional profiles and marker genes for diverse supraspinal populations. J Neurosci:JN-RM-1197-22 .

Blesch A, Uy HS, Grill RJ, Cheng JG, Patterson PH, Tuszynski MH (1999) Leukemia inhibitory factor augments neurotrophin expression and corticospinal axon growth after adult CNS injury. J Neurosci 19:3556–3566 .

Cafferty WBJ, Gardiner NJ, Das P, Qiu J, McMahon SB, Thompson SWN (2004) Conditioning injury-induced spinal axon regeneration fails in interleukin-6 knock-out mice. J Neurosci 24:4432–4443 .

Campbell IL, Abraham CR, Masliah E, Kemper P, Inglis JD, Oldstone MBA, Mucke L (1993) Neurologic disease induced in transgenic mice by cerebral overexpression of interleukin 6. Proc Natl Acad Sci U S A 90:10061–10065 .

Cao Z, Gao Y, Bryson JB, Hou J, Chaudhry N, Siddiq M, Martinez J, Spencer T, Carmel J, Hart RB, Filbin MT (2006) The cytokine interleukin-6 is sufficient but not necessary to mimic the peripheral conditioning lesion effect on axonal growth. J Neurosci 26:5565– 5573 .

Cui Q, Harvey AR (2000) CNTF promotes the regrowth of retinal ganglion cell axons into murine peripheral nerve grafts. Neuroreport 11:3999–4002 .

Fischer D (2017) Hyper-IL-6: a potent and efficacious stimulator of RGC regeneration. Eye (Lond) 31:173–178 .

Fischer M, Goldschmitt J, Peschel C, Brakenhoff JPG, Kallen KJ, Wollmer A, Grötzinger J, Rose-John S (1997) I. A bioactive designer cytokine for human hematopoietic progenitor cell expansion. Nat Biotechnol 15:142–145 .

Geoffroy CG, Meves JM, Kim HJM, Romaus-Sanjurjo D, Sutherland TC, Li JJ, Suen J, Sanchez JJ, Zheng B (2022) Targeting PTEN but not SOCS3 resists an age-dependent decline in promoting axon sprouting. iScience 25:105383 .

He Z, Jin Y (2016) Intrinsic Control of Axon Regeneration. Neuron 90:437–451 .

Hodgetts SI, Yoon JH, Fogliani A, Akinpelu EA, Baron-Heeris D, Houwers IGJ, Wheeler LPG, Majda BT, Santhakumar S, Lovett SJ, Duce E, Pollett MA, Wiseman TM, Fehily B, Harvey AR (2018) Cortical AAV-CNTF Gene Therapy Combined with Intraspinal Mesenchymal Precursor Cell Transplantation Promotes Functional and Morphological Outcomes after Spinal Cord Injury in Adult Rats. Neural Plast 2018 .

Imai Y, Kohsaka S (2002) Intracellular signaling in M-CSF-induced microglia activation: role of Iba1. Glia 40:164–174.

Jin D, Liu Y, Sun F, Wang X, Liu X, He Z (2015) Restoration of skilled locomotion by sprouting corticospinal axons induced by co-deletion of PTEN and SOCS3. Nat Commun 6:8074 Available at: http://www.ncbi.nlm.nih.gov/pubmed/26598325.

Lang C, Bradley PM, Jacobi A, Kerschensteiner M, Bareyre FM (2013) STAT3 promotes corticospinal remodelling and functional recovery after spinal cord injury. EMBO Rep 14:931–937 Available at: http://www.ncbi.nlm.nih.gov/pubmed/23928811 [Accessed May 19, 2017].

Leaver SG, Cui Q, Plant GW, Arulpragasam A, Hisheh S, Verhaagen J, Harvey AR (2006a) AAV-mediated expression of CNTF promotes long-term survival and regeneration of adult rat retinal ganglion cells. Gene Ther 13:1328–1341.

Leaver SG, Cui Q, Plant GW, Arulpragasam A, Hisheh S, Verhaagen J, Harvey AR (2006b) AAV-mediated expression of CNTF promotes long-term survival and regeneration of adult rat retinal ganglion cells. Gene Ther 13:1328–1341.

Leibinger M, Andreadaki A, Gobrecht P, Levin E, Diekmann H, Fischer D (2016) Boosting central nervous system axon regeneration by circumventing limitations of natural cytokine signaling. Molecular Therapy 24:1712–1725.

Leibinger M, Müller A, Gobrecht P, Diekmann H, Andreadaki A, Fischer D (2013) Interleukin-6 contributes to CNS axon regeneration upon inflammatory stimulation. Cell Death Dis 4.

Leibinger M, Zeitler C, Gobrecht P, Andreadaki A, Gisselmann G, Fischer D (2021) Transneuronal delivery of hyper-interleukin-6 enables functional recovery after severe spinal cord injury in mice. Nat Commun 12:1–14.

Mahar M, Cavalli V (2018) Intrinsic mechanisms of neuronal axon regeneration. Nat Rev Neurosci 19:323–337.

O’Donovan KJ (2016) Intrinsic Axonal Growth and the Drive for Regeneration. Front Neurosci 10:486 Available at: http://www.ncbi.nlm.nih.gov/pubmed/27833527

Rose-John S (2012) IL-6 trans-signaling via the soluble IL-6 receptor: importance for the pro-inflammatory activities of IL-6. Int J Biol Sci 8:1237–1247.

Rose-John S (2018) Interleukin-6 Family Cytokines. Cold Spring Harb Perspect Biol 10:a028415

Sahenk Z, Seharaseyon J, Mendell JR (1994) CNTF potentiates peripheral nerve regeneration. Brain Res 655:246–250

Smith PD, Sun F, Park KK, Cai B, Wang C, Kuwako K, Martinez-Carrasco I, Connolly L, He Z (2009) SOCS3 Deletion Promotes Optic Nerve Regeneration In Vivo. Neuron 64:617–623

Sun F, Park KK, Belin S, Wang D, Lu T, Chen G, Zhang K, Yeung C, Feng G, Yankner BA, He Z (2011) Sustained axon regeneration induced by co-deletion of PTEN and SOCS3. Nature 480:372–375

Tedeschi A, Bradke F (2017) Spatial and temporal arrangement of neuronal intrinsic and extrinsic mechanisms controlling axon regeneration. Curr Opin Neurobiol 42:118–127

Vartak A, Goyal D, Kumar H (2023) Role of Axon Guidance Molecules in Ascending and Descending Paths in Spinal Cord Regeneration. Neuroscience 533:36–52

Wang Z, Kumaran M, Batsel E, Testor-Cabrera S, Beine Z, Alvarez Ribelles A, Tsoulfas P, Venkatesh I, Blackmore MG (2025) Single-nuclei sequencing reveals a robust corticospinal response to nearby axotomy but overall insensitivity to spinal injury. J Neurosci:e1508242024

Wang Z, Maunze B, Wang Y, Tsoulfas P, Blackmore MG (2018) Global connectivity and function of descending spinal input revealed by 3D microscopy and retrograde transduction. The Journal of Neuroscience:1196–18

Wang Z, Romanski A, Mehra V, Wang Y, Brannigan M, Campbell BC, Petsko GA, Tsoulfas P, Blackmore MG (2022) Brain-wide analysis of the supraspinal connectome reveals anatomical correlates to functional recovery after spinal injury. Elife 11

Yang P, Qin Y, Bian C, Zhao Y, Zhang W (2015) Intrathecal Delivery of IL-6 Reactivates the Intrinsic Growth Capacity of Pyramidal Cells in the Sensorimotor Cortex after Spinal Cord Injury. PLoS One 10:e0127772

Yang P, Wen H, Ou S, Cui J, Fan D (2012) IL-6 promotes regeneration and functional recovery after cortical spinal tract injury by reactivating intrinsic growth program of neurons and enhancing synapse formation. Exp Neurol 236:19–27

Zhong J, Dietzel ID, Wahle P, Kopf M, Heumann R (1999) Sensory impairments and delayed regeneration of sensory axons in interleukin-6-deficient mice. J Neurosci 19:4305–4313

Zigmond RE (2012) Cytokines That Promote Nerve Regeneration. Exp Neurol 238:101

